# Crosstalk between Nitric Oxide and Retinoic Acid pathways is essential for amphioxus pharynx development

**DOI:** 10.1101/2020.06.22.164632

**Authors:** F Caccavale, G Annona, L Subirana, H Escriva, S Bertrand, S D’Aniello

## Abstract

During animal ontogenesis, body axis patterning is finely regulated by complex interactions between several signaling pathways. Nitric Oxide (NO) and Retinoic Acid (RA) are potent morphogens that play a pivotal role in vertebrate development. Their involvement in axial patterning of head and pharynx shows conserved features in the chordate phylum. Indeed, in the cephalochordate amphioxus NO and RA are crucial for the correct development of pharyngeal structures. Here we demonstrate the functional cooperation between NO and RA occurring in amphioxus embryogenesis. During neurulation, NO modulates RA production through the transcriptional regulation of Aldh1a.2 that irreversibly converts retinaldehyde into RA. On the other hand, RA regulates the transcription of Nos genes, probably through RA Response Elements found in their regulatory regions. The reciprocal regulation of NO and RA pathways results to be essential for the normal pharyngeal development in amphioxus and suggests that this regulatory crosstalk could be conserved in vertebrates.

## Introduction

The ontogenesis of the vertebrate head is a complex developmental process in which both neural crest and non-neural crest cells participate. The craniofacial development and the correct antero-posterior patterning of head structures are driven by complex interactions between several signaling pathways and epigenetic mechanisms (Haworth et al., 2007; Jacox et al., 2014; Kong et al., 2014; Francis-West and Crespo-Enriquez, 2016). In this context Nitric Oxide (NO) is a potent morphogen playing crucial roles in head structures development. Loss-of-function of neuronal Nitric Oxide Synthase (*Nos1*) in *Xenopus* and zebrafish induces the arrest of mouth opening, smaller eye and strong aberrations in cartilage and bone structures formation (Jacox et al., 2014). Moreover, the inhibition of NO production is responsible for severe defects in pharyngeal arches patterning, consistent with alterations in the Hox code (Kong et al., 2014).

The development and the antero-posterior and dorso-ventral patterning of the head and pharynx show conserved features within the chordate phylum. In amphioxus, which belongs to the cephalochordate subphylum, pharynx is characterized by a marked left-right asymmetry which is controlled by the Nodal signaling pathway, namely by the Cerberus-Nodal-Lefty-Pitx cascade (Bertrand et al., 2015; Soukup et al., 2015; Li et al., 2017). The antero-posterior patterning and development of amphioxus pharyngeal slits are driven by a conserved set of transcription factor genes, among which *Hox1, Pax2/5/8, Pitx, Tbx1/10*, and *Engrailed* (Schubert et al., 2005; Bertrand et al., 2015; Wang et al., 2019) similarly to the development of pharyngeal arches in vertebrates.

NO is enzymatically produced in amphioxus by three distinct NO synthase genes: *NosA, NosB* and *NosC*, derived from cephalochordate-specific gene duplications and showing a complementary expression pattern during development (Annona et al., 2017). During amphioxus embryonic development, as a consequence of pharmacological inhibition of endogenous NO production, the opening of the mouth is prevented as well as the correct development of other important pharyngeal structures, such as the endostyle and the club-shaped gland (Annona et al., 2017). Moreover, the larvae show a posteriorized phenotype, resembling the well described phenotype induced by RA administration during amphioxus embryogenesis (Escriva et al., 2002; Schubert et al., 2005; Koop et al., 2014). These experimental evidences prompted us to investigate the hypothesis of a possible evolutionarily conserved role of NO and RA in chordate pharynx development using the cephalochordate *Branchiostoma lanceolatum* as a model system. In order to highlight the molecular mechanisms driving developmental changes in the amphioxus embryo, we took advantage of the transcriptomic differences induced by pharmacological treatments reducing endogenous NO production, which alter the pharyngeal area development (Annona et al., 2017). Our approach allowed us to demonstrate that such morphogenetic alterations are linked to a dramatic imbalance affecting the reciprocal regulation of NO and RA pathways.

## Results

### NO controls pharyngeal development during early neurulation in amphioxus

Previous studies have highlighted the involvement of NO in the specification of amphioxus pharyngeal structures (Annona et al., 2017). To better characterize the key role of NO during embryonic development, we decided to narrow down the time window of treatment by defining the exact timing during which NO is functional. Therefore, we performed short-term *in vivo* treatments with the NOS activity inhibitor 1-[2-(trifluoromethyl)phenyl]-1H-imidazole (TRIM) during *B. lanceolatum* development using different drug exposure times between early neurula stage (N2 stage) and pre-mouth larva stage (transition stage T2) (Figure 1A). The resulting phenotype was then observed at the open-mouth larva stage (larva-L0 stage). The observed morphological alterations included: *i*. a reduced antero-posterior length of the pharynx, *ii*. a complete or partial absence of mouth opening on the left side of the pharynx (Figure 1B, section c and c’), *iii*. an incomplete formation of the club-shaped gland and of the endostyle (Figure 1B, section d, e and d’, e’), and *iv*. a smaller first gill slit (Figure 1B, section f and f’).

**Figure 1.**
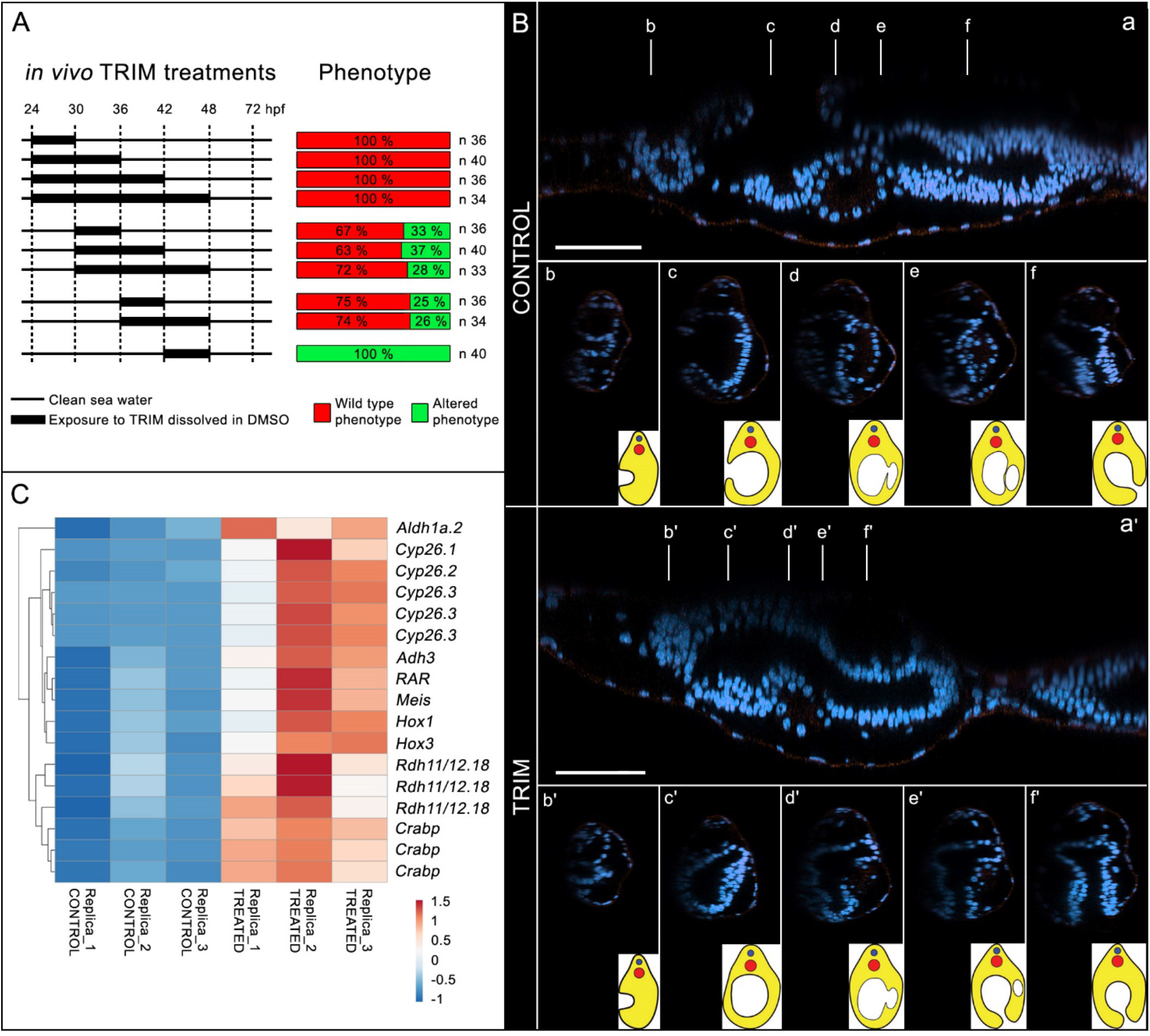
Phenotypic characterization of in vivo TRIM treatment during early amphioxus embryogenesis. A) Schematic representation of time intervals during which embryos were grown in presence of TRIM and the obtained resulting phenotype. B) Digital sections of control and TRIM-treated larvae after DAPI staining showing alteration in pharyngeal region. Anterior to the left. C) Gene expression heatmap of differential transcriptomic analysis (control versus TRIM).

The inhibition of NO production during the time window 24-30 hours post fertilization (hpf, at 18°C) was the shortest treatment inducing a strong effect by producing 100% of abnormal larval while delayed treatments starting at 30 or 36 hpf, for 6, 12 or 24 hours, resulted in approximately 70% of affected larvae. Conversely, when TRIM treatment was performed later, between 42 and 48 hpf, the larvae were not affected (Figure 1A). These results suggest, therefore, that pharyngeal development is, at least in part, under the control of NO during neurulation and that the critical time of its action is restricted approximately to six hours. Based on these experimental evidences, we performed a differential transcriptomic analysis comparing TRIM-treated embryos (continuous treatment from 24 to 30 hpf) with wild-type embryos in order to define the genes acting downstream of NO signaling during pharyngeal development in amphioxus (Figure. 1C - figure supplement 1).

### Inhibition of NO synthesis *in vivo* induces up-regulation and ectopic expression of RA pathway genes

The differential RNA-Seq analysis of TRIM-treated embryos (30 hpf) revealed the up-regulation of 392 and the down-regulation of 50 genes (Figure 1 C - figure supplement 1 panels A-B-C). Interestingly, several differentially up-regulated genes are implicated in RA metabolism and signaling pathway (synthesis and storage, catabolism, and known direct RA target genes): *Adh3, Rdh11/12.18, Aldh1a.2, Crabp, Cyp26.1, Cyp26.2, Cyp26.3, RAR, Hox1, Hox3, Meis* (Figure 1C – figure supplement 2). In order to confirm this finding, the expression levels of several of these genes were additionally analyzed by quantitative RT-PCR (qRT-PCR). The results showed a consistent expression change trend with the RNA-seq data (Figure 1C - figure supplement 1 panel D). Moreover, the expression pattern of RA target genes *Hox1, Hox3, Meis* and *Cyp26* genes was further investigated by whole-mount *in situ* hybridization in both control and TRIM-treated embryos at the neurula N4 and pre-mouth T2 developmental stages. Such analyses showed that endogenous NO reduction produced an effect not only on the expression level of RA metabolism and signaling pathway genes, but also on the expression territories of most of them. *Hox1, Hox3* and *Meis* anterior limit of expression was pushed anteriorly in TRIM-treated embryos in comparison to controls, indicative of the embryo posteriorization (Figure 2A). The RA catabolism genes that are duplicated in amphioxus, *Cyp26.1, Cyp26.2, Cyp26.3*, denoted an heterogeneous behavior: *Cyp26.2* was slightly up-regulated and its expression pattern did not change after TRIM treatment, whereas *Cyp26.1* and *Cyp26.3* were strongly up-regulated and showed an ectopic expression (Figure 2A). In particular, after inhibition of NO production, *Cyp26.1* expression was anteriorized, while *Cyp26.3* expression was posteriorized. Moreover, *Cyp26.3* showed an additional domain of expression in the tailbud, mainly in the pre-mouth larvae (Figure 2A).

**Figure 2.**
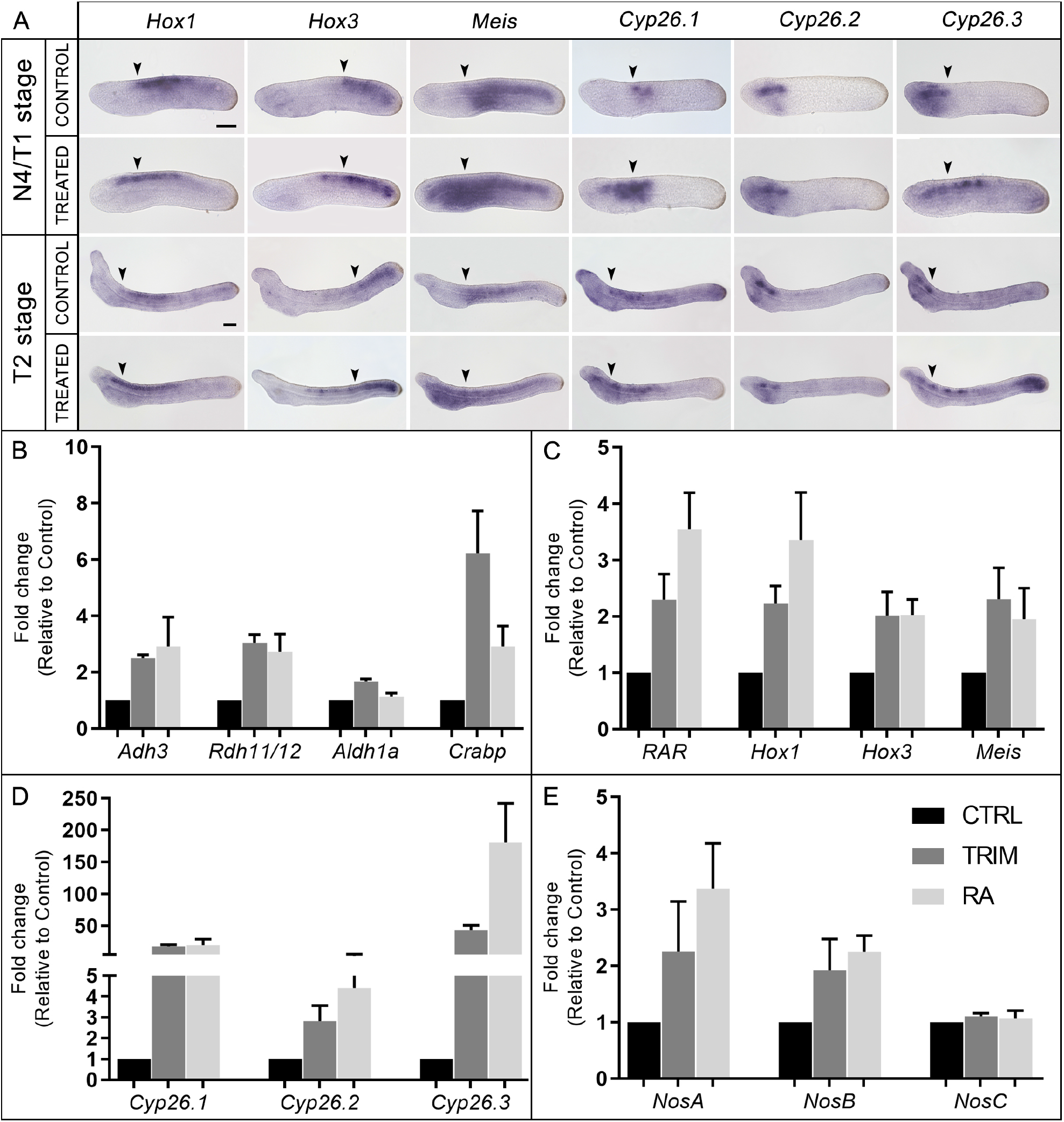
Analysis of gene expression. **A)** Gene expression pattern by *in situ* hybridization of *Hox1, Hox3, Meis, Cyp26.1, Cyp26.2* and *Cyp26.3* after the inhibition of NO production and in controls in 30 hpf (N4/T1 stage) and 42 hpf (T2) embryos. The anterior (*Hox1, Hox3, Meis, Cyp26.1*) and posterior (*Cyp26.3*) limits of gene expression territories in a wild type embryo are indicated with arrowheads in both control and TRIM-treated conditions showing the posteriorization (or anteriorization for *Cyp26.3*) of gene expression in treated embryos. Embryos orientation: anterior to the left, dorsal to the top. Scale bars: N4/T1 stage: 50μM; T2 stage: 50μM. **B-E)** Histograms of qRT-PCR experiment results show expression level changes, after 6 hours of pharmacological TRIM or RA treatments, of: **B)** genes encoding enzymes for RA synthesis: *Adh3, Rdh11/12.18, Aldh1a.2* and binding protein for storage: *Crabp;* **C)** genes known as direct targets of RA: *Hox1, Hox3, Meis, RAR;* **D)** genes encoding RA degradation enzymes: *Cyp26.1, Cyp26.2, Cyp26.3;* **E)** *Nos* genes: *NosA, NosB, NosC*. The gene expression analyses were performed by qRT-PCR on 30 hpf embryos (N4) RNA samples. The statistical analysis showed a pvalue <0,05 for all the data, except for *NosA* after TRIM in **(E)** that show a pvalue = 0,07.

### *Aldh1a.2* is specifically regulated by NO

The above-mentioned gene expression results suggested that the abnormal pharyngeal development of TRIM-treated embryos could be the result of an up-regulation of the RA signaling pathway. In order to test this hypothesis, we performed two *in vivo* experiments in parallel in which early neurula embryos were incubated for 6 hours, from 24 to 30 hpf, in the presence of either TRIM or RA. Then, we analyzed the relative expression of three groups of genes, that previously we found to be up-regulated in our RNA-seq analysis, by qRT-PCR: *i*. genes involved in the synthesis and in storage of RA, as *Adh3, Rdh11/12, Aldh1a.2, Crabp; ii*. genes that mediate RA effects, as *RAR, Hox1, Hox3, Meis;* and *iii*) genes involved in RA degradation, as *Cyp26.1, Cyp26.2, Cyp26.3*. All analyzed genes were up-regulated after both TRIM and RA treatment, with the exception of *Aldh1a.2* which was exclusively up-regulated after TRIM treatment (Figure 2 B-C-D).

### *NosA* and *NosB* respond to exogenous RA during development

The expression analysis of the three amphioxus *Nos* genes after TRIM treatment revealed the transcriptional up-regulation for two of them, *NosA* and *NosB*, while *NosC*, the only one constitutively expressed during development (Annona et al., 2017) remained insensitive to the pharmacological treatment (Figure 2E).

In order to check if *NosA* and *NosB* up-regulation could be due to an indirect effect of the intracellular increase of RA caused by the TRIM treatment, we tested their expression levels after the addition of exogenous RA (Figure 2E). Similarly, to the TRIM treatment, we observed that *NosA* and *NosB* expression were significantly up-regulated also as a consequence of RA administration. Moreover, exogenous RA induces up-regulation of *NosA* expression up to 36 hpf and *NosB* expression up to 48 hpf (Figure 2E - figure supplement 3 panel A).

These experimental evidences suggest a possible transcriptional effect of RA on *Nos* genes during embryogenesis in amphioxus. To further support this assumption, we searched for the presence of Retinoic Acid Response Elements (RARE) in the putative regulatory genomic regions of *NosA* and *NosB*, using a computational prediction tool. We looked for RAREs in selected open chromatin regions identified by the overlapping peaks obtained from ATAC-seq and Chip-seq data (Marlétaz et al., 2018). Six putative Direct Repeat (DR)-type binding sites were detected in the *NosA* locus (two DR1, three DR3 and one DR5), and only one DR5 in the 5’ region of *NosB* (Figure supplement 3 panel B), suggesting a possible direct RA regulation of *Nos* genes in amphioxus.

### A RALDH inhibitor and a RAR antagonist are able to rescue the normal phenotype after inhibition of NO synthesis

To confirm that the up-regulation of *Aldh1a.2*, which could result in an endogenous RA increase, is the key event underlying pharyngeal alterations in TRIM-treated larvae, we performed two independent phenotypic rescue experiments using the RALDH inhibitor DEAB (N,N-diethylaminobenzaldehyde) and the RA antagonist BMS009. Both DEAB and BMS009 were applied in combination with TRIM to embryos at 24 hpf and removed at 30 hpf. As a control, the TRIM treatment was performed in parallel on another batch of embryos. The combined treatment with TRIM and DEAB resulted in the recovery of the wild type phenotype in 76% of the total observed larvae, while 14% of them showed a partially recovered phenotype with a smaller mouth compared to controls (Figure 3A, B). The rescue experiments performed using the combination of TRIM and BMS009 led to 54% of complete and 21% of partial recovery of the wild type larval morphology (Figure 3A). Moreover, these morphological rescue experiments were associated with the recovery at N4 neurula stage of the normal expression pattern of RA catabolism (*Cyp26.1* and *Cyp26.3*) and RA target (*Hox1, Hox3* and *Meis*) genes (Figure 3C).

**Figure 3.**
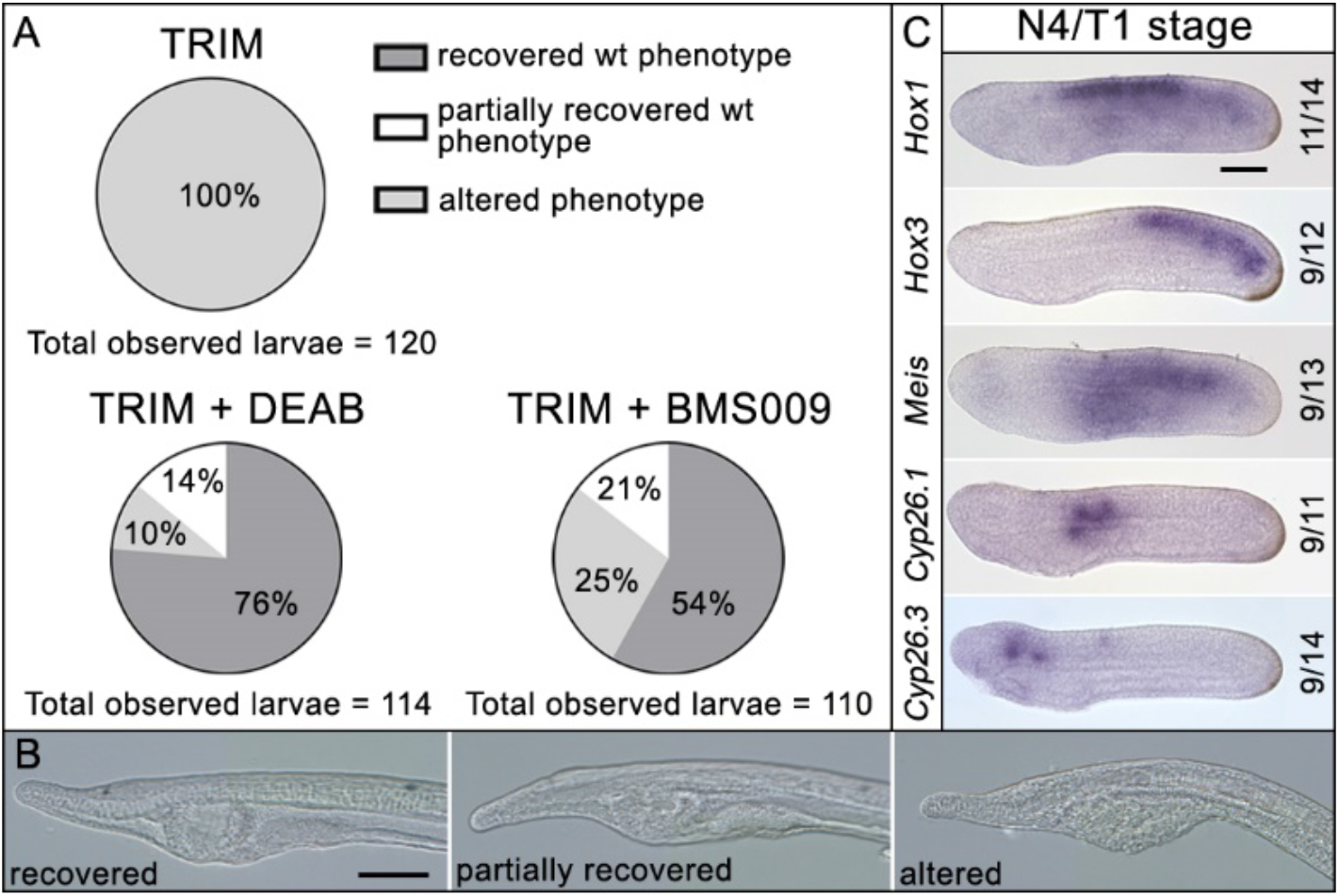
Phenotypic rescue effect of DEAB and BMS009 on TRIM-treated embryos. A) Pie charts of the phenotypes observed after TRIM treatment and the combinatorial pharmacological treatments TRIM (100 μM) + DEAB (25 μM) and TRIM (100 μM) + BMS009 (10-6 M). The percentages of each observed phenotype are reported in the respective portions of the graphs. For each treatment, the total number of observed larvae is indicated below the chart. B) Pictures of the pharyngeal region of larvae presenting the three different classes of phenotype observed in the rescue experiments: recovered wild type phenotype, partially recovered wild type phenotype, altered phenotype. C) Gene expression pattern analyses by *in situ* hybridization of *Hox1, Hox3*, Meis, *Cyp26.1*, and *Cyp26.3* after rescue assay with DEAB show the restoration of wild-type gene expression territories of the analyzed genes. *Cyp26.2* expression was not assayed because its localization in the embryo does not change after TRIM treatment. The numbers indicate the ratio between embryos showing a restored expression pattern and the total number of embryos analyzed. Embryos orientation: anterior to the left, dorsal to the top. Scale bar: 50 μM.

Therefore, by using two independent experiments, we demonstrate that the reduction of RA levels, or of its regulatory action, in TRIM-treated embryos, rescues the wild type phenotype, suggesting that the observed phenotype in TRIM-treated embryos is produced by an increase in RA signaling.

## Discussion

### NO controls RA synthesis by the transcriptional regulation of *Aldh1a.2*

The inhibition of NO production during amphioxus neurulation affects the normal formation and localization of pharyngeal structures at the larva stage, including the length of the pharynx itself. From a molecular point of view, the differential RNA-seq approach revealed a clear up-regulation of different RA pathway players after Nos activity inhibition, suggesting that such de-regulation is responsible for the observed phenotype. The role of RA in pharyngeal morphogenesis has been extensively described in the literature; RA acts through *Hox1* in establishing the posterior limit of amphioxus pharynx. *Hox1* is co-expressed with the RA receptor (RAR) in the midgut endoderm and, in turn, represses the expression of pharyngeal endoderm markers, such as *Pax1/9* and *Otx* (Schubert et al., 2005). Nevertheless, the formation of pharyngeal slits requires low levels of RA. This condition is guaranteed *i*. by the activity of RA degradation enzymes (Cyp26), *ii*. by the expression of TR2/4, a transcriptional repressor which binds on RAREs and decreases RA signaling in the anterior part of the animal, and *iii*. by the fact that the embryonic region producing RA (the central region of the embryo) moves posteriorly as the embryo elongates (Escriva et al., 2002; Koop et al., 2014). In our studies, as a result of Nos activity inhibition, the RA target genes, *Hox1, Hox3* and *Meis*, showed an increased expression level and an anterior limit of expression which is moved anteriorly, consistent with the posteriorization of the larva body plan. Furthermore, the RA degrading enzyme genes, *Cyp26.1* and *Cyp26.3*, were also sensible to the inhibition of endogenous NO production showing an increased and ectopic expression. *Cyp26* genes are required for RA degradation in endoderm and ectoderm and have a key role in the establishment and maintenance of the antero-posterior RA concentration gradient in amphioxus (Carvalho et al., 2017a). The up-regulation of Cyp26 genes is a known consequence of RA excess, which is responsible of the posteriorization of larval body structures and of the pharynx loss (Escriva et al., 2002; Schubert et al., 2005; Schubert et al., 2006; Minoux and Rijli, 2010; Koop et al., 2010, 2014; Bertrand et al., 2015; Carvalho et al., 2017b). Altogether, these results suggested that the observed phenotype in the TRIM-treated amphioxus embryos could be due to an increase in RA production. Intracellular RA is synthetized by the reversible oxidation of retinol in retinaldehyde by either alcohol dehydrogenases (ADH) or retinol dehydrogenases (RDH). Subsequently, retinaldehyde is irreversibly oxidized to RA by retinaldehyde dehydrogenases (RALDH), mainly by ALDH1A (Gallego et al., 2006; Duester, 2008). In particular, in our experiments we observed transcriptional up-regulation of *Adh3, Rdh11/12.18* and *Aldh1a.2* after TRIM treatments. The unique orthologue of vertebrate *Adh* genes, *Adh3*, was identified in amphioxus (Cañestro et al., 2002). Instead, 22 *Rdh11/12* genes in amphioxus, derived from a lineage-specific expansion, were identified as related to human *Rdh11, Rdh12, Rdh13* and *Rdh14*, corresponding to retinaldehyde reductases predominantly involved in retinoid metabolism and homeostasis (Albalat et al., 2011) (Figure supplement 2). For *Aldh1*, a total of six genes were identified in amphioxus, orthologs of human *Aldh1A1-3*, which are major players in the oxidation of RA (Cañestro et al., 2006) (Figure supplement 2). In our study, *Adh3* and *Rdh11/12.18* genes also resulted up-regulated after administration of exogenous RA, suggesting a feedback regulation of RA synthesis by itself, at least on reversible enzymatic steps. On the other hand, *Aldh1a.2* was insensible to exogenous RA excess. Based on these results we hypothesized that, under physiological conditions, NO transcriptionally regulates *Aldh1a.2* and, as a consequence, controls the production of endogenous RA. Further evidence to support our hypothesis is provided by two independent rescue experiments. Technically, in association with TRIM, we used two drugs that specifically act on most crucial steps of RA signaling pathway: an inhibitor of the RALDH enzymes activity that would compensate the excess of Aldh1a.2, and a RA antagonist able to compensate the RA over-production through its binding to RAR. In both rescue experiments we observed not only the recovery of the wild type phenotype but also the restoration of the expression pattern of both RA-target and RA-degrading genes.

Therefore, based on our results, we propose NO as a key player in the fine regulation of RA synthesis during neurulation in amphioxus. Thus, NO fine-tune the expression of *Aldh1a.2*, keeping RA concentration within the optimal range. This precise balance between intracellular concentrations of NO and RA guarantees the correct expression levels and territory localization of all RA downstream target genes. When NO is removed from the system the RA metabolism machinery malfunctions, giving rise to a consequent cascade of events that lead to the up-regulation of the entire RA signaling pathway.

The missing piece of the puzzle, therefore, seems to be an unknown protein able to mediate the control of *Aldh1a.2* transcription by NO. The activity of such protein could depend on its phosphorylation and S-nitrosylation state that would be modulated by NO. Indeed, in other chordates, it has been demonstrated that the mechanisms by which NO regulates transcription of target genes are: *i*. the control of the extracellular-regulated kinase (ERK) and the MAP kinase phosphatases (MAPK) activity and, as a result, the modulation of phosphorylation or dephosphorylation of target transcription factors (Castellano et al., 2014), and *ii*. the direct modulation of target proteins, as transcription factors, histone acetyltransferases and deacetylases or DNA methyltranserases, through S-nitrosylation of specific cysteine residues (Bogdan, 2001; Nott et al., 2008; Sha and Marshall, 2012).

Hence, further research is necessary to find out which is the NO-target protein that mediates *Aldh1a.2* expression regulation and how NO specifically controls its activity. It could be important improve this knowledge since very little information, and only restricted to vertebrates, is reported on the control of RA metabolism by NO. Some cytochrome P450 enzymes, involved in RA–metabolism, were identified as putative NO–regulated proteins but no evidences about such a possible transcriptional regulation have been reported (Lee et al., 2014, 2017).

### RA controls NO synthesis

The exogenous administration of RA induces the expression of two amphioxus *Nos* genes that are normally not expressed during development, *NosA* and *NosB* (Annona et al., 2017). Moreover, this activation is maintained throughout the whole critical time period during which NO is necessary for pharyngeal development (Figure supplement 3 panel A). Conversely, *NosC*, the only *Nos* gene expressed during embryogenesis in normal conditions, seems not to be affected by the increase of RA level.

The *in silico* analysis of the chromosomal loci of both *B. lanceolatum NosA* and *NosB* revealed the presence of seven DR-type RAREs (Figure supplement 3 panel B), localized in open chromatin regions detected by ATAC-seq and Chip-seq approaches. This suggests a possible regulation of *Nos* genes by RA through the binding of RAR/RXR nuclear receptors to these putative RAREs. The transcriptional activation of *NosA* and *NosB* by RA during development could be part of a mechanism to rebalance the correct NO/RA ratio. However, this hypothesis should be further confirmed by experimental validation.

In vertebrates, the role of NO and RA in the correct development of the pharynx and craniofacial structures has already been described. For example, NO is described to be necessary for face formation as part of the Kinin-Kallikrein pathway (Jacox et al., 2014) and, on the other hand, the correct RA gradient, generated by local degradation, leads to appropriate specification of craniofacial structures (Abe et al., 2008; Liu et al., 2013; Chawla et al., 2018).

However, a possible crosstalk between NO and RA pathways has not yet been revealed. As mentioned above, there are few published data on the regulatory effect of NO on RA pathway, while the control of RA on NO production in vertebrates is more documented. For instance, in human cells, it has been demonstrated the inhibitory or activation effect of RA on NO production through activation of inducible or endothelial NOS at both protein and transcript levels (Sirsjö et al., 2000; Achan et al., 2002; Hattori et al., 2002; Behairi et al., 2015; Moon, 2019).

## Conclusions

Our results show the presence of a functional crosstalk between RA and NO signals in the cephalochordate amphioxus during neurulation. This opens new questions about the evolutionary conservation of this regulatory loop in the development of the pharynx and the head of vertebrates.

The role of RA as well as that of NO in amphioxus development and antero-posterior patterning of the pharynx has been described in several studies. Our results suggest the occurrence of a regulatory crosstalk between these two ancient and essential signaling pathways that has been previously neglected. Based on our data we propose that, during amphioxus development, a precise NO/RA balance is necessary for the correct antero-posterior patterning of the pharynx. This endogenous intracellular balance is preserved by the reciprocal regulation of NO and RA pathways. Given such evidences and knowledge from vertebrate literature, we hypothesize that the functional and regulatory crosstalk between NO and RA pathways could be a conserved feature in vertebrates.

## Materials & Methods

### Amphioxus embryos collection

Ripe adult European amphioxus (*Branchiostoma lanceolatum*) were collected in Argelès-sur-mer (France) with a specific permission delivered by the Prefect of Region Provence Alpes Côte d’Azur. *B. lanceolatum* is not a protected species. Spawning was induced during late spring and beginning of summer by employing a thermal shock as described by Fuentes *et al*. (Fuentes et al., 2007) After *in vitro* fertilization, embryos were cultured in 0,22 μm filtered seawater at 18°C in plastic Petri dishes. Following the staging at 18°C, 24 hpf corresponds to neurula 2 stage (N2), 30 hpf to neurula 4 stage (N4), 36 hpf to transition 1 stage (T1), 42 hpf to transition 2 stage (T2), 48 hpf to transition 3 stage (T3), and 72 hpf to larva 0 (L0). At desired developmental stages embryos were used for specific drug treatments, and then they were frozen in liquid nitrogen and kept at −80°C for subsequent RNA extraction, or fixed with 4% paraformaldehyde (PFA) in MOPS buffer overnight at 4°C and then stored in 70% ethanol at - 20°C.

### Pharmacological treatments

Amphioxus embryos at different developmental stages were treated with the NOS inhibitor 1-[2-(trifluoromethyl)phenyl]-1H-imidazole (TRIM), with the RALDH inhibitor N,N-diethylaminobenzaldehyde (DEAB), with the RA antagonist BMS009 and with Retinoic Acid (RA). All the drugs were dissolved in dimethyl sulfoxide (DMSO), and control embryo groups for each treatment were prepared adding an equal amount of DMSO. For TRIM treatments, a final concentration of 100 μM was used starting at 24 hpf. Embryos at 30, 36, 42 and 48 hpf were collected or were rinsed in filtered seawater and allowed to develop until 72 hpf to observe phenotype. For RA treatments, a final concentration of 10^-6^ M was used. Embryos were collected at the developmental stages of 30, 36, 42 and 48 hpf. All the treatments were performed in biological triplicates.

For rescue experiments, embryos at 24 hpf were treated simultaneously with a combination of 100 μM TRIM and 25 μM DEAB, or 100 μM TRIM and 10^-6^ M BMS009. At 30 hpf they were rinsed in filtered seawater and allowed to develop until 72 hpf stage when the phenotype was observed. The experiment was performed in biological duplicates.

Control and TRIM-treated larvae at 72 hpf were stained with DAPI. A high-resolution Z-stack was acquired using a Zeiss confocal microscopy LSM 800, and a medium speed fast interactive deconvolution was applied. For the digital sections, Imaris 9.3.1 software was used.

### RNA-seq analysis

Total RNA was extracted from embryos using the RNeasy Plus Mini Kit (QIAGEN) after sample homogenization using the TissueLyser (QIAGEN). The RNA integrity number (RIN) was assessed by using TapeStatio4200 while RNA concentration and purity were estimated using a Nanodrop spectrophotometer. Indexed libraries were prepared from 1 ug/ea purified RNA with TruSeq Stranded Total RNA Library Prep Kit. Libraries were quantified using the Agilent 2100 Bioanalyzer (Agilent Technologies) and pooled so that each index-tagged sample was present in equimolar amounts, with a final concentration of 2 nM. The pooled samples at a final concentration of 10 pM were subjected to cluster generation and sequencing using an Illumina NextSeq500 System in a 1×75 single read format (30 millions of reads). The raw sequence files generated (fastq files) underwent quality control analysis using FastQC. Transcriptome sequences were deposited in the NCBI Sequence Read Archive (SRA) database with the accession number: PRJNA630453.

Reads were mapped on the *B. lanceolatum* transcriptome (Oulion et al., 2012) using the aligner Bowtie2 with default parameters (Langmead and Salzberg, 2012). The read counts were obtained using IdxStats (Li et al., 2009; Cock et al., 2013) and the differential expression analysis between treated and wild-type embryos was performed using the R package DESeq2 (Love et al., 2014). Mapping and read counting were performed on the Roscoff ABiMS galaxy platform.

### Phylogenetic analysis

Phylogenetic analysis was necessary to establish the orthology relationships of the RA synthesis genes differentially expressed in our experiments. Protein alignments were generated with ClustalX program using the sequence database reported in Handberg-Thorsager *et al*. (Handberg-Thorsager et al., 2018). Phylogenetic trees were based on maximum-likelihood inferences calculated with PhyML v3.0 (Guindon et al., 2010) (Figure supplement 2).

### Computational prediction of DR-type RAREs in *NosA* and *NosB* genomic loci

We selected open chromatin genomic regions in the vicinity of *Nos* genes by choosing regions corresponding to ATAC-seq peaks (8, 15, 36, 60 hpf) overlapping with ChiP-seq signals for the H3K4me3 mark at the same developmental times. The prediction of putative Direct Repeats (DR) binding sites in the genomic loci of *NosA* and *NosB* was assessed by NHR-SCAN tool (Sandelin and Wasserman, 2005), using the following parameter: 0,01 combined probability of entering match states.

### Gene expression analysis by whole-mount *in situ* hybridization

*Hox1, Hox3, Meis, Cyp26.1, Cyp26.2, Cyp26.3* were cloned in pGEM-T vector (Promega) using primers listed in Supplementary table 1. Antisense riboprobes were synthesized and *in situ* hybridizations were performed as previously described (Annona et al., 2017). Embryos were mounted in 80% glycerol in PBS, and photographed using an Axio Imager.Z2.

### Gene expression analysis by qRT-PCR

Total RNA was extracted from embryos at different developmental stages: 30, 36, 42 and 48 hpf (Figure 1A), using the RNeasy Plus Mini Kit (QIAGEN). 350-1000 ng of total RNA were retrotranscribed in cDNA which was used undiluted (only for *Nos* genes) or diluted 1:10 for the quantitative PCR. Each qPCR reaction contained a final concentration of 0.7 μM of each primer and Fast SYBR Green Master mix with ROX (Applied Biosystems) in 10 μl total volume. PCR reactions were run in a ViiA™ 7 Real-Time PCR System (Applied Biosystems). The cycling conditions were: 95°C for 20 s, 40 cycles with 95°C for 1 s, 60°C for 20 s, 95°C for 15 s, 60°C 1 min, followed by a dissociation curve analysis using a gradient from 60°C to 95°C with a continuous detection at 0.015°C/sec increment for 15 min. The results were analyzed using the ViiA™-7 Software and exported into Microsoft Excel for further analysis. Each sample was processed in biological triplicates. Ribosomal protein L32 (*Rpl32*), expressed at a constant level during development, was used as a reference gene for the normalization of each gene expression level (Annona et al., 2017). Primers used are listed in Supplementary table 1. For the statistical analysis, we used the GraphPad Prism software employing the T-test. Statistical significance cut off criteria was set at p<0.05.

## Acknowledgments

The authors thank Rebecca Adikes and Hannah Rosenblatt, course assistants at the MBL Embryology Course 2019 in Woods Hole (USA), for their help with imaging. We are also grateful to Enrico D’Aniello and Ricard Albalat for their critical reading of the manuscript. We are grateful to the Institut Français de Bioinformatique and the Roscoff Bioinformatics platform ABiMS for providing computing and storage resources for the RNA-seq analysis. We thank the BIO2MAR platform for giving us access to analytical material.

SD and FC acknowledge the Assemble Plus Project (contract numbers BA010618 and 360BA0619) and The Company of Biologists (grant number: DEVTF-170211; Sponsoring Journal: Development) for supporting research visits to the Observatoire Océanologique of Banyuls-sur-Mer (France). FC was supported by an OU-SZN PhD fellowship. SB is supported by the Institut Universitaire de France (IUF). HE is supported by the Centre national de la recherche scientifique and Agence Nationale de la Recherche (ANR) grants no. ANR-16-CE12-0008-01 and ANR-19-CE13-0011.

---

**Figure supplement 1.**
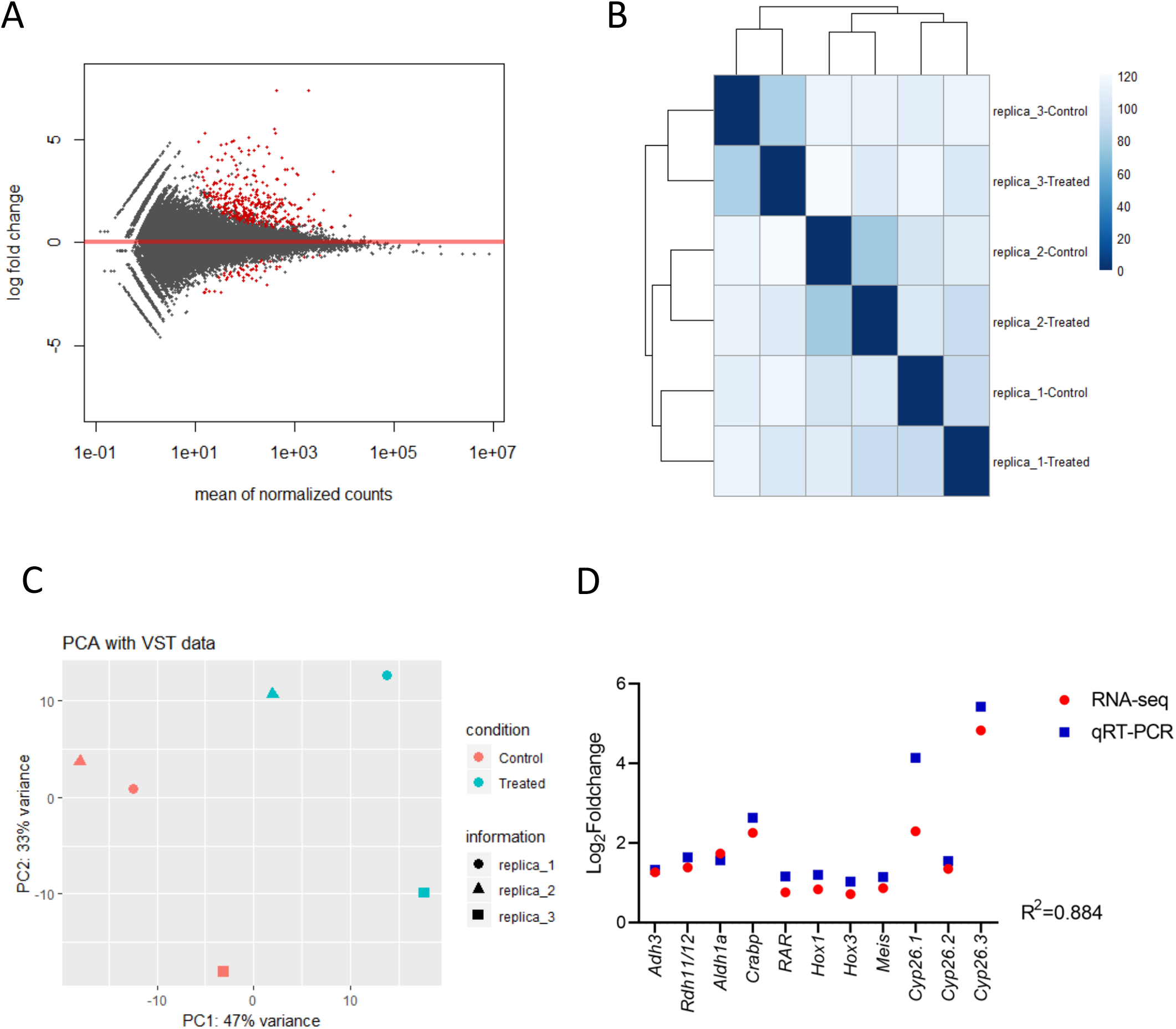
Quality of RNA-seq data. A) MA plot, red dots plotted represent genes with an adjusted *p* value <0,1 (treated vs control). B) Heatmap of sample-to-sample distances, strains are clustered by replicas (3 controls and 3 treated); C. Principal component analysis (PCA) plot. Sample classes are highlighted in different colors: control in red and treated in blue. D) Gene expression correlation between RT-qPCR and RNA-seq data for the following selected genes: Adh3, Rdh11/12.18, Aldh1a.2, Crabp, RAR, Hox1, Hox3, Meis, Cyp26.1, Cyp26.2 and Cyp26.3. Blu squares indicate Log2 of relative foldchanges obtained from RT-qPCR analysis of TRIM-treated versus untreated samples. Red circles indicate Log2 of relative foldchanges obtained from RNAseq analysis of TRIM-treated versus untreated samples. The Pearson correlation Coefficient (R2) is indicated.

**Figure supplement 2.**
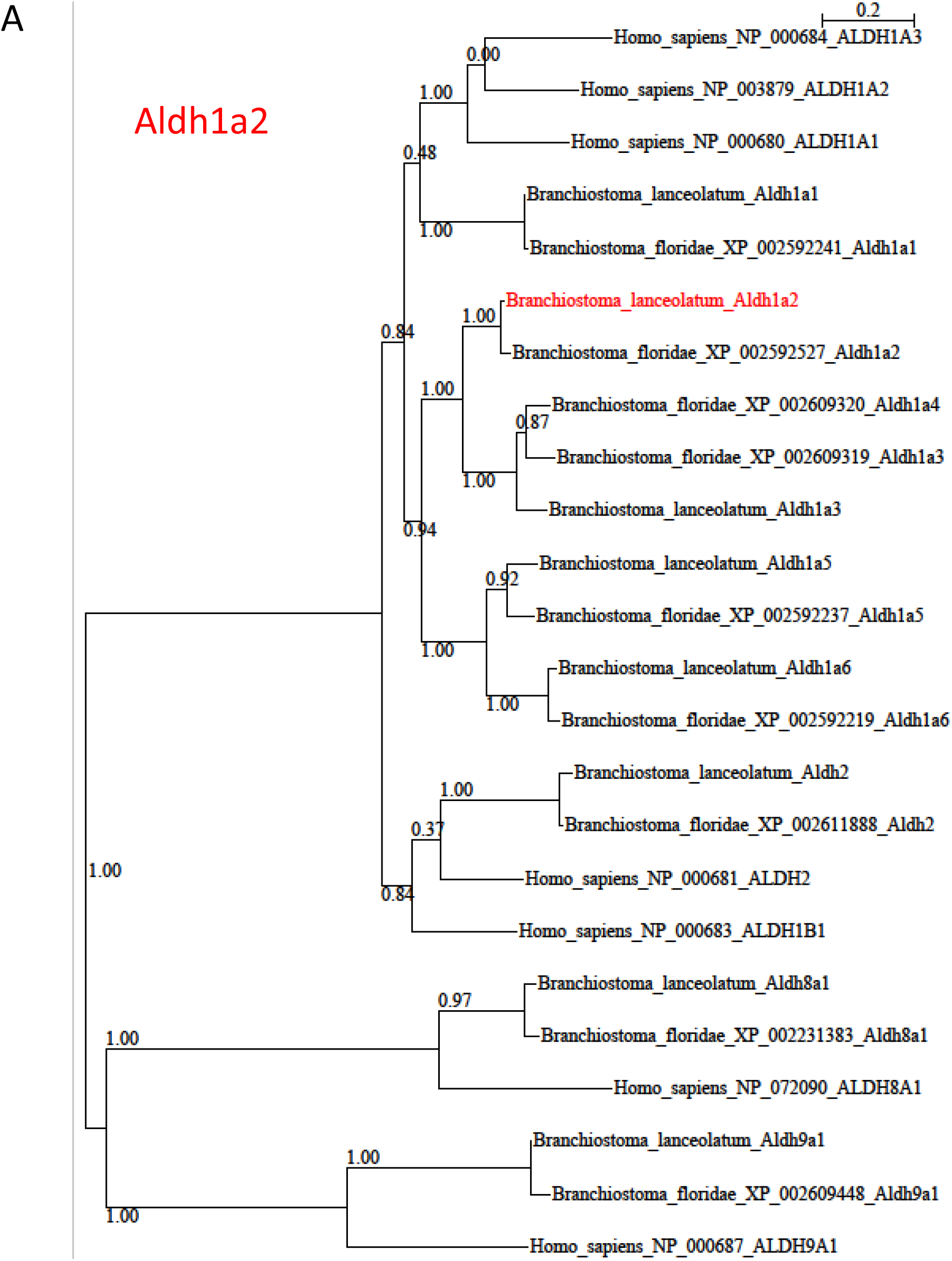
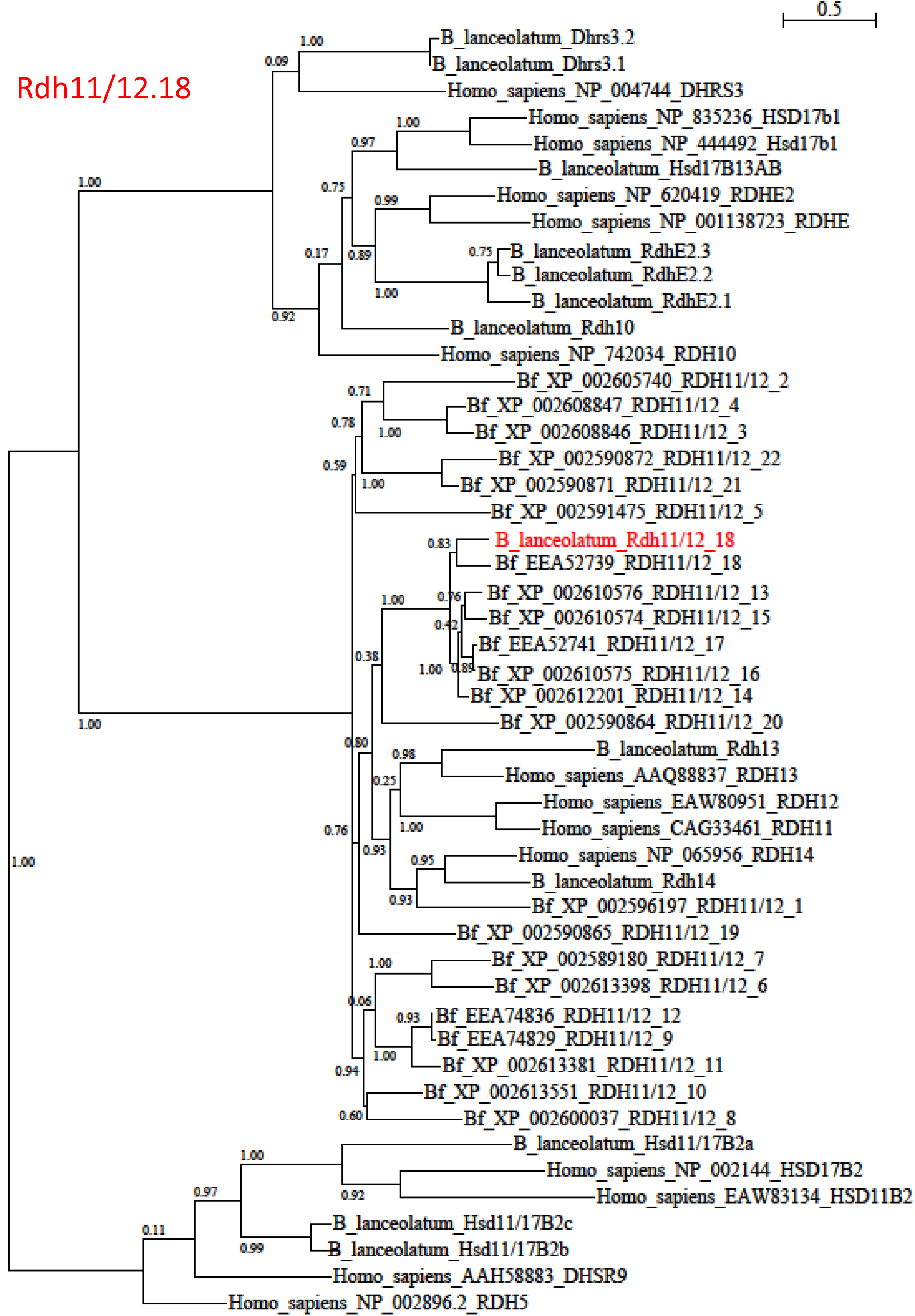
Phylogenetic analysis of RA pathway genes resulted upregulated in the differential WT versus TRIM transcriptome. The analysis allowed the identification of *Branchiostoma lanceolatum* homologs (indicated in red in the trees) for (A) Aldh1a2 (ContigAmph4820) and (B) Rdh11/12.18 (ContigAmph8913/ContigAmph35515/ContigAmph81467). Trees were calculated using Maximum Likelihood (ML) method, and bootstrap supports are given at each node. The GenBank accession number is indicated for each protein sequence used.

**Figure supplement 3.**
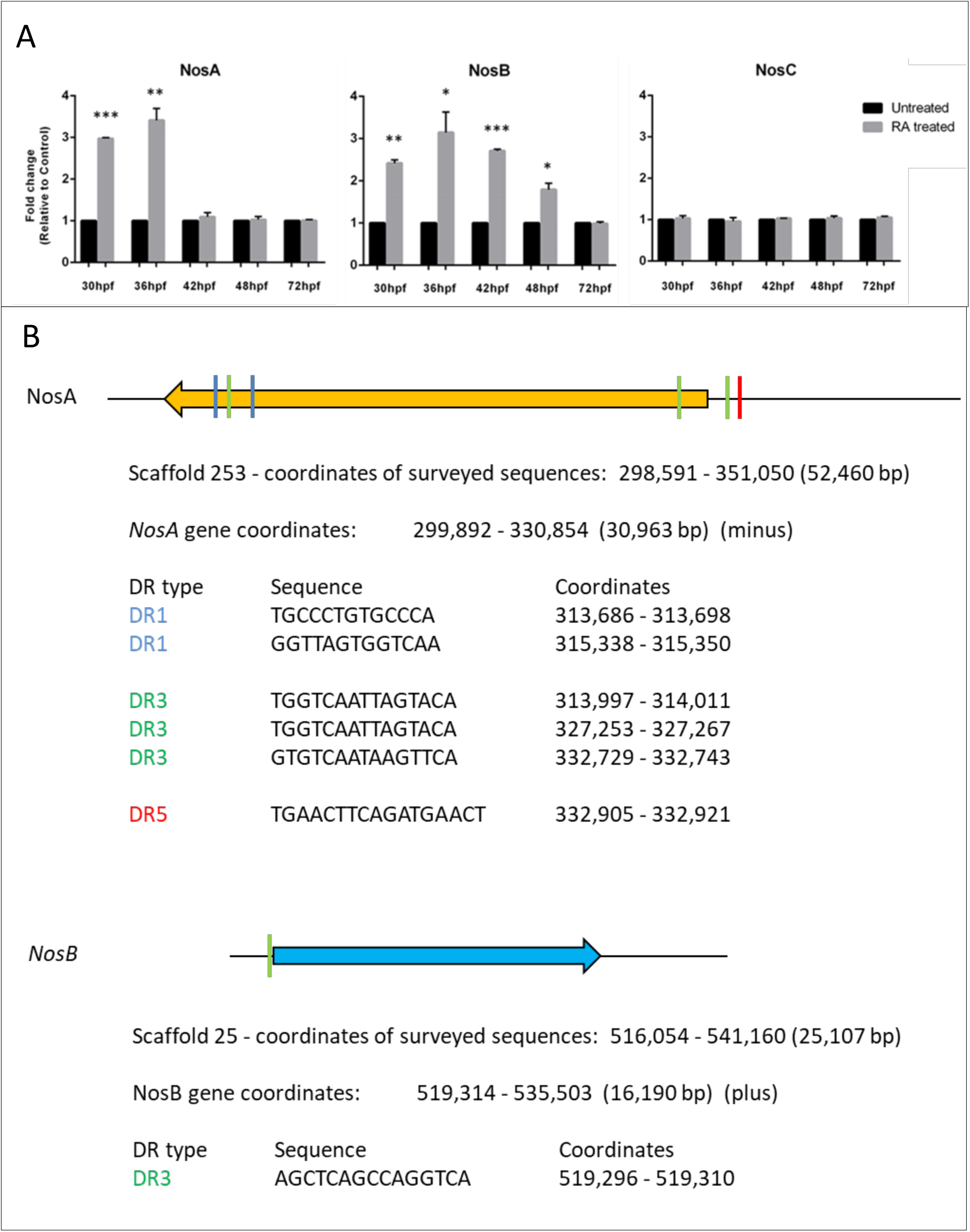
**A)** *Nos* genes expression pattern by RT-qPCR after RA treatment during amphioxus development. Five developmental time points were assayed from 30 to 72 hours post-fertilization. * = pvalue<0,05; **= pvalue<0,005; ***= pvalue<0,001. **B)** *In silico* prediction analysis of RARE elements in genomic loci of Branchiostoma lanceolatum NosA and NosB. The analysis was limited to neighboring genes on both sides of each Nos gene. Direct Repeat 1 (DR1) are shown in blue, DR3 in green and DR5 in red. Sequence and coordinates of each putative DR are indicated.

**Supplementary table 1.**
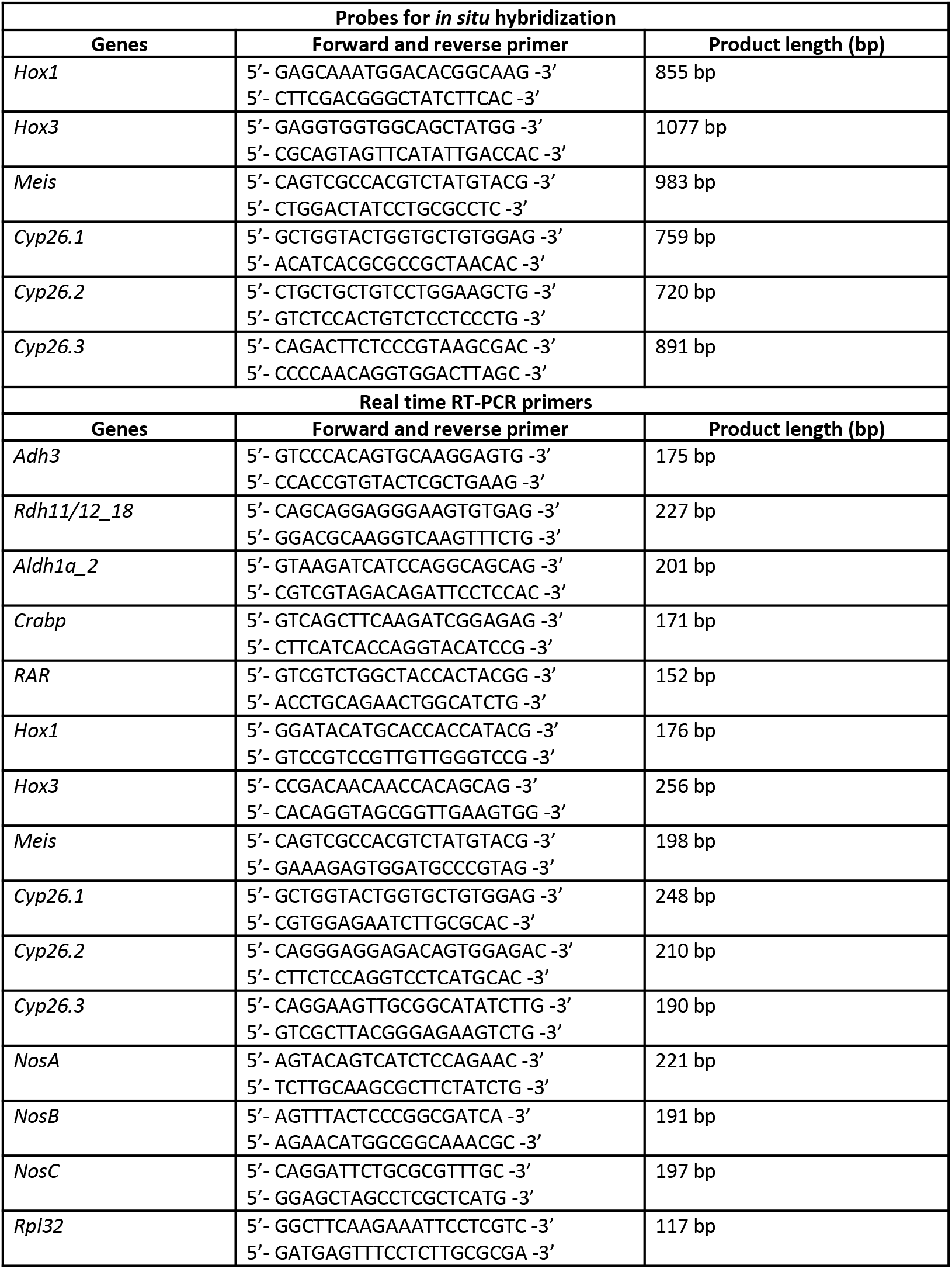
Oligonucleotides used to amplify probes for *in situ* hybridization experiments and for Real time RT-PCR analyses.

